# Older adults generalize their movements across walking contexts more than young during gradual and abrupt split-belt walking

**DOI:** 10.1101/2021.08.06.455403

**Authors:** Yashar Aucie, Harrison. M. Harket, Carly. J. Sombric, Gelsy Torres-Oviedo

**Affiliations:** Sensory Motor Learning Laboratory, University of Pittsburgh, Department of Bioengineering, Pittsburgh, PA, USA

**Keywords:** motor learning, rehabilitation robotics, gait, aging, kinematics

## Abstract

Generalization of movements from experienced to novel situations is a critical aspect of motor learning. It has been demonstrated that the training period when a movement is learned influences the movement’s generalization to untrained situations. However, little is known about how healthy aging affects these processes. For example, young adults exhibit greater generalization of movements learned on a device (e.g. split-belt treadmill or robotic arm) to movements without it (e.g, overground walking or unconstrained reaching) when participants experience small vs. large perturbations on the training device. Here, we investigated whether a similar effect would be observed in older adults. To this end, we compared the generalization of split-belt adaptation to overground walking in older (75.9±4.8 years old) and young adults (24.7±5.9 years old) when adapted gradually (i.e., small perturbations) vs. abruptly (i.e., large perturbations). We found that both age groups adapted more to the abrupt condition compared to the gradual condition, which resulted in greater adaptation effects (i.e., aftereffects) on the treadmill in the abrupt than the gradual groups. We also found that older adults generalize more than young adults, regardless of the perturbation schedule (i.e., gradual or abrupt). Our results suggest that abrupt perturbations during adaptation do not limit the generalization of movement in older adults-perhaps because they are more likely to attribute them to their own faulty movements. These results suggest that large perturbations are better than small when training older clinical populations since abrupt disturbances would lead to more adaptation and generalization of corrected movements in older people.

## 1 Introduction

As the world’s population grows older (Ortman et al., 2014), understanding how mechanisms of motor adaptation change with healthy aging, and how to counteract these age-related changes, becomes increasingly more important. Healthy older adults wish to maintain an independent (and active) lifestyle despite changes in their bodies or their surroundings (Nelson et al., 2007). For older adults to continue performing daily activities, they must maintain a flexible motor system that counteracts endogenous or exogenous perturbations to their movements through motor adaptation (King et al., 2013). Thus, it is important to understand age-related changes in motor adaptation. It has been shown that healthy aging impairs the rate of adaptation (Fernández-Ruiz et al., 2000; Sombric et al., 2017) and the extent of adapted movements (McNay and Willingham, 1998) when a novel environment is suddenly experienced. This seems to be a general trait of the aged motor system as it has been observed during reaching (Buch et al., 2003; Bock, 2005) and walking movements (Bruijn et al., 2012; Sombric et al., 2017). Age-related decline in adaptation performance could be attributed to impaired cognitive strategies. Specifically, older adults have difficulties identifying the external disturbances altering their movements (Buch et al., 2003; Bock, 2005; Heuer and Hegele, 2008; Hegele and Heuer, 2010), challenging their ability to consciously counteract them (Hegele and Heuer, 2013; Vandevoorde and de Xivry, 2020). On the other hand, there is no consensus on the impact of healthy aging on implicit processes underlying sensorimotor adaptation (Mazzoni and Krakauer, 2006; Sülzenbrück and Heuer, 2009; Taylor and Ivry, 2011), some suggest that this is preserved with healthy aging (Heuer and Hegele, 2008; Vandevoorde and de Xivry, 2020), whereas some others have shown age related decline in implicit motor adaptation (Wolpe et al., 2016; Iturralde and Torres-Oviedo, 2019). Thus, we investigated the extent to which small perturbations recruiting implicit processes (Rommech2015) would enhance the motor adaptation capacity in older individuals.

The generalization of adapted movements from training to testing contexts is an important aspect of motor adaptation. Namely, generalization is defined as the ability to carry over information from trained experiences to novel situations (Krakauer, 2009; Torres-Oviedo et al., 2011). For example, an expert tennis player can generalize their motor repertoire to learn faster table tennis than someone without tennis experience. Thus, one can exploit generalization to benefit motor performances in new situations. This generalization capacity is critical for the efficacy of robotic-assisted rehabilitation. Namely, if devices like exoskeletons or treadmills are to be used as training devices, patients must generalize the movements learned on the devices to real-life situations without them. Therefore, there is an interest in finding factors facilitating the generalization of corrected movements beyond the clinical setting. Previous studies have shown that older adults generalize their movements more than young individuals (Fernández-Ruiz et al., 2000; Sombric et al., 2017; Sombric and Torres-Oviedo, 2021). However, the adaptation effects (i.e., aftereffects) remain significantly larger in the training environment (e.g., treadmill walking) compared to those in a different environment (e.g., regular overground walking) (Reisman et al., 2009; Sombric et al., 2017; Sombric and Torres-Oviedo, 2021). This raises the question of whether the adaptation experience in older adults could be manipulated to increase the generalization of adapted movements. For example, small perturbations (i.e., gradual adaptation) of reaching (Kluzik et al., 2008) and walking (Torres-Oviedo and Bastian, 2012) in young adults and post-stroke individuals (Alcântara et al., 2018) result in a larger generalization of movements. It is, however, unknown whether a similar effect can be observed in older adults.

In this study, we investigated the extent to which small vs. large perturbations can regulate the generalization of locomotor adaptation in young and older adults. We hypothesized that small perturbations (i.e., gradual adaptation) would lead to more generalization in both age groups compared to large perturbations (i.e., abrupt adaptation). To test this hypothesis, young and older adults adapted their walking pattern on a split-belt treadmill either gradually or abruptly. We compared the adaptation and generalization of movements across these groups. Interestingly, older adults generalized more than young, regardless of the perturbation schedule, suggesting that healthy aging reduces the ability to contextualize motor memories in older populations.

## 2 Methods

### 2.1 Participants

We investigated if adaptation experience could be manipulated to increase the generalization of movements from the split-belt treadmill to overground walking in older adults. To this end, we adapted 32 healthy adults on a split-belt treadmill either gradually (i.e., small perturbations) or abruptly (i.e., large perturbations). Sixteen older (10 males and 6 females, mean age 75.9±4.8 years old) and sixteen young participants (8 males and 8 females, mean age 24.7±5.9 years old) experienced an abrupt or gradual perturbation. Older and young adults were randomly assigned to either perturbation group, yielding four groups (i.e., Old_Abrupt_, Old_Gradual,_ Young_Abrupt_, and Young_Gradual_) of 8 participants each. Sixteen additional older adults (10 males and 6 females, mean age 76±5 years) were collected in a post-hoc experiment. These individuals were adapted gradually (Old_gradual_NC_) or abruptly (Old_abrupt_NC_) without a catch condition in which the adaptation period was briefly interrupted to measure aftereffects on the treadmill. These additional groups were tested to ensure that our observations were not due to the unintended large perturbations that the gradual groups experienced after said catch condition (see detailed explanation in the Statistical Analysis section of this article). Participants did not have sensory, neurological, or musculoskeletal disorders. The Institutional Review Board at the University of Pittsburgh approved the experimental protocol and all participants gave informed consent before testing.

### 2.2 Locomotor Paradigm

All participants walked both overground and on a treadmill during the experiment to complete a conventional generalization protocol that consisted of three walking epochs: baseline, adaptation, and post-adaptation (Figure 1A-top). First, participants experienced the baseline epoch overground, during which they walked back-and-forth on a 9-meter walkway (i.e., overground walking) for 6 minutes (~ 150 strides) before walking on the treadmill. Then, participants experienced the baseline epoch on the treadmill, during which they walked on the treadmill at three different speeds, which were slow (0.5 m/s), fast (1 m/s), and medium (0.75 m/s) speeds for 150 strides each. Next, participants experienced either an abrupt or gradual split-belt adaptation epoch for 600 strides. For the abrupt groups only, participants took a short (~ 3 min) break after every 150 strides to allow older participants to rest. Participants in the gradual group did not take breaks because we wanted to avoid the errors that older adults experience after a break (Sombric 2017). Besides, gradual adaptation did not require as many breaks because it was less strenuous than abrupt adaptation. Speed profiles for each error size are shown in Figure 1A-bottom. In the abrupt case, the belts suddenly moved at a 2:1 belt ratio (1 and 0.5m/s), whereas in the gradual case, one belt sped up from 0.75 to 1m/s as the other one slowed down from 0.75 to 0.5 m/s. The faster belt was under the dominant leg for every participant, which was determined by self-report of preferred kicking leg (Kramer and Balsor, 1990). A brief catch condition (10 strides) during which both belts moved at 0.75 m/s was used to assess treadmill aftereffects. We chose this speed because it is approximately the effective speed at which participants walk during the adaptation epoch. Following the catch condition, participants were re-adapted to the split-belt perturbation for 300 strides (before walking overground). Lastly, in the post-adaptation epoch, participants walked overground, immediately after the re-adaptation period, for 6 minutes to assess the generalization of treadmill aftereffects to a different walking condition. Participants did not take any transition steps between walking on the treadmill and walking overground. Following the overground walking, participants walked again on the treadmill for 300 strides when the two belts moved at 0.75 m/s to assess the remaining aftereffects that were specific to the treadmill context.

**Figure 1.**
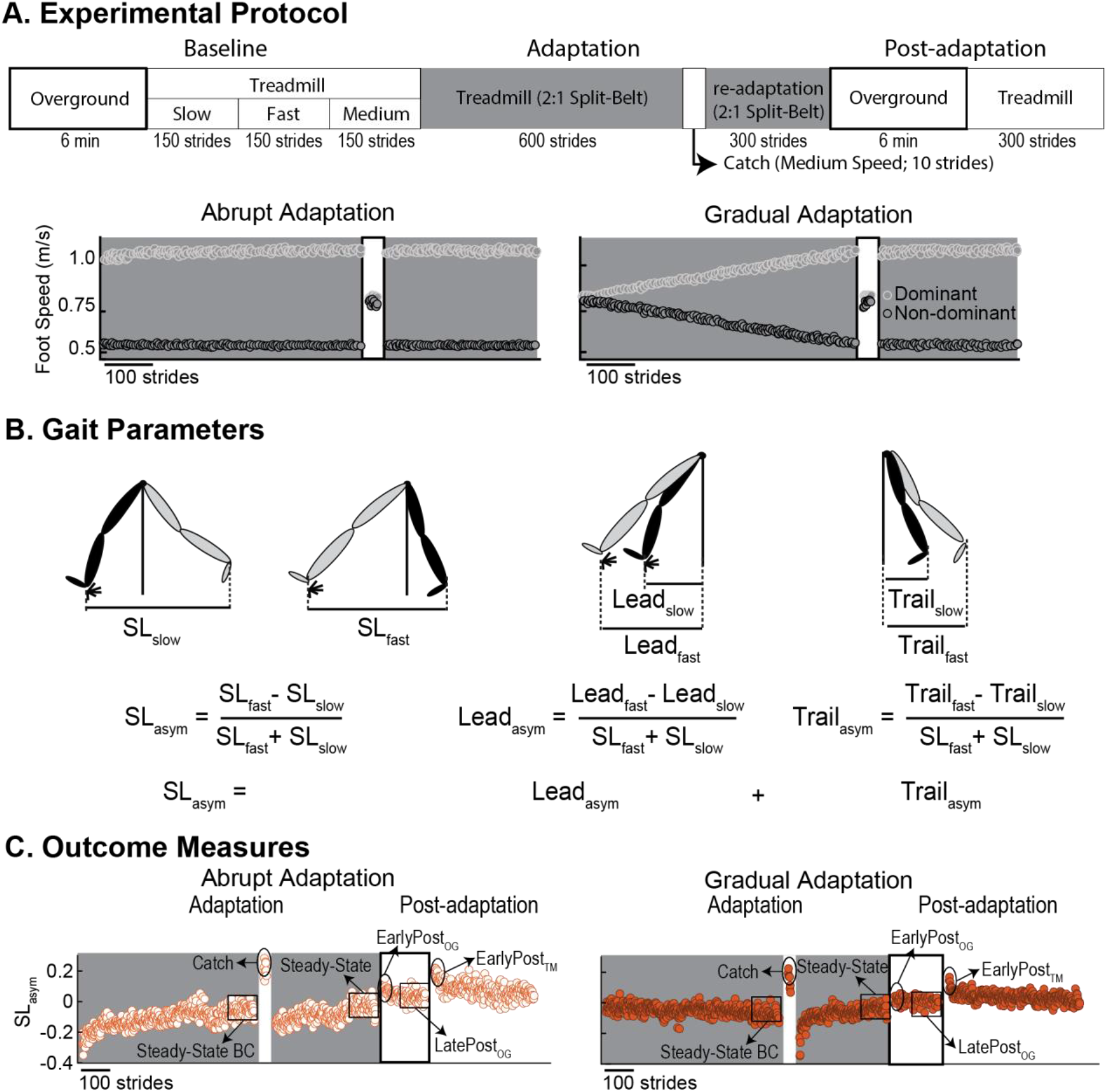
(A-top) Experimental protocol. The paradigm used for all groups consisted of 3 phases: baseline, adaptation, and post-adaptation. Thick black outlines represent overground walking. Thin black outlines represent treadmill conditions with both belts moving at the same speed (i.e., tied). The gray background represents the adaptation and re-adaptation periods during which the dominant leg is moving two times faster than the non-dominant leg (i.e., split). The labels “Slow”, “Fast”, and “Medium” refer to the speeds at which both legs were moving: 0.5 m/s, 1m/s, and 0.75/s, respectively. (A-bottom) Speed profiles during adaptation. The graph illustrates the time course of the foot speed at which the dominant (white circles) and non-dominant legs (black circles) walked during the adaptation period. (B) This schematic illustrates step length asymmetry and its decomposition into leading and trailing leg asymmetries. Step length asymmetry is quantified as the difference between fast and slow step lengths, normalized by stride length. The equation and decomposition are explained in detail in the section “Materials and Methods” of this article. In brief, the asymmetry between the fast (gray) and the slow (black) leading leg’s positions contribute to step length asymmetry. Similarly, asymmetry in the trailing leg’s positions also contributes to step length asymmetry. (C) Outcome measures. Epochs of interest are illustrated over an example step length asymmetry time course. The shaded gray area represents the adaptation period when the feet move at different speeds (“split” walking), whereas white areas represent when the feet move at the same speed. The thick black outline represents the overground condition.

For safety purposes, all individuals wore a ceiling-mounted harness during the entire paradigm that only provided support in the event of a fall. For the treadmill walking conditions, participants were alerted when the treadmill was about to start and stop, but were not informed about the speed of the belts. Participants were instructed to hold on to a handrail positioned in front of them at the beginning and end of each treadmill condition, but were encouraged to let go as soon as they felt comfortable walking with their arms unrestricted (as they did during overground walking). Participants were also instructed to look straight ahead while walking so that they would not be distracted by the motion of the belts, which has been shown to alter the generalization of movements (Mariscal et al., 2020). An examiner stood by to monitor compliance with these instructions. Also, a plastic divider was placed between the treadmill belts to ensure that participants could not step on the wrong belt when walking on the treadmill.

### 2.3 Data Collection

Kinematic and kinetic data were collected at 100 Hz and 1000 Hz, respectively, using a passive motion capture system (Vicon Motion Systems, Oxford UK), and an instrumented split-belt treadmill (Bertec, Columbus OH). Positions from the ankle (lateral malleolus) and the hip (greater trochanter) were collected bilaterally. Markers were also placed asymmetrically on the shanks and thighs to differentiate between the legs. Gaps in raw kinematic data due to marker occlusion were filled by visual inspection of each participant using the Vicon Nexus software. Ground reaction forces recorded by force plates under each treadmill belt were used to count in real-time the number of strides that participants walked on the treadmill. Following data collection, instances of heel-strikes (i.e. foot landing) and toe-offs (i.e., foot lifting) were identified using kinematic data. This was done to have equivalent event detection on the treadmill and overground as in previous generalization studies (Torres-Oviedo and Bastian, 2010, 2012; Sombric et al., 2017; Mariscal et al., 2020; Sombric and Torres-Oviedo, 2021). Custom MATLAB scripts were used to perform all data analysis.

### 2.4 Data Analysis

#### Gait Parameters

We characterized the adaptation and generalization patterns of every group using step length asymmetry, which is a metric conventionally used to quantify adaptation and generalization of gait in split-belt protocols. Step length asymmetry (SL_asym_) was defined as the difference between step lengths (SL, the distance between ankles) when taking a step with the leg walking slow vs. the leg walking fast (Eq. 1) (Figure 1B). SL_fast_ is defined as the distance between ankles at fast heel strike (i.e., Fast leg is leading) and vice versa for SL_slow_ (i.e., Slow leg is leading). A zero value of step length asymmetry indicated that both step lengths were equal and a positive value indicated that the step length of the fast (dominant) leg was longer than the slow (non-dominant) leg. We further decomposed step length asymmetry into asymmetries between the leading (Lead_asym_) or trailing (*Trail*_*asym*_) positions (Figure 1B) because these have been shown to generalize differently (Mariscal et al. 2020). *Lead*_*asym*_ (Eq. 2) and *Trail*_*asym*_ (Eq. 3) were calculated as follows:

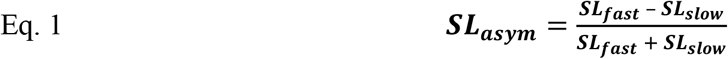

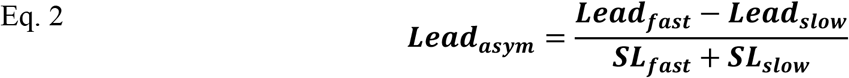

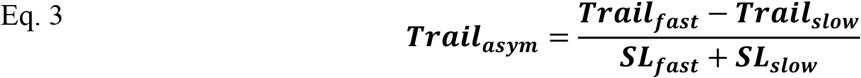

In these equations, the leading leg’s position (Lead_fast_ or Lead_slow_) was defined as the ankle’s marker position in the sagittal plane of the leg in front of the body at heel-strike and the trailing leg’s position (Trail_fast_ or Trail_slow_) was defined as that of the leg behind the body. Both positions are computed with respect to the body, which is defined as the averaged position of the two hip markers at heel-strike. Moreover, the leading leg’s position *Lead*_*fast*_ was the position of the fast leg when this was in front of the body at fast heel-strike, whereas *Lead*_*slow*_ was the same but when the slow leg was the leading leg. By convention, positive *Lead*_*asym*_ values indicated that the fast leg landed farther forward from the body compared to when the slow leg landed forward when taking a step. Similarly, the trailing leg’s position *Trail*_*fast*_ was the position of the fast leg when this was the trailing leg at slow heel strike and vice versa for *Trail*_*slow*_. By convention, large negative *Trail*_*asym*_ values indicated that the fast leg was farther behind the body compared to when the slow leg was behind the body at the contralateral heel strike.

#### Outcome Measures

One outcome measure was maximum error size. This was computed as the average of the first 5 steps of step length asymmetry, SL_asym_, measured when the 2 belts were at their maximum speed difference (i.e., Full split). The full split occurs at the beginning of the adaptation period for the abrupt groups, whereas it happens at the beginning of the re-adaptation period (after the catch) for the gradual groups. We quantified the maximum error size by averaging the SL_asym_ during the first 5 strides of the full split period. We chose SL_asym_ as a global measure of error size since this is a performance metric that is robustly minimized as people adapt during split-belt adaptation paradigms (Reisman et al., 2005; Finley et al., 2015).

Six other outcome measures were computed for each gait parameter at specific epochs of interest within the experimental protocol. These outcome measures consisted of 1) steady-state before the catch (**SteadyStateBC**), 2) aftereffects during the catch (**Catch**), 3) steady-state before overground walking (**SteadyState**), 4) early aftereffects overground (**EarlyPost**_**OG**_), 5) late aftereffects overground (**LatePost**_**OG**_), and 6) remaining aftereffects on the treadmill (**EarlyPost**_**TM**_). These outcome measures were used to compare the adaptation and generalization between the groups (Figure 1C). In all outcome measures, we first removed the five strides at the end of each epoch to eliminate the effect of slowing down the treadmill’s belts before stopping.

We quantified the steady-state before catch (**SteadyStateBC**, average of last 40 strides) to contrast the adapted states across groups before measuring the aftereffects on the treadmill during the catch. Next, we quantified the aftereffects during the catch period (**Catch**, average of the first 5 strides) to assess the aftereffects on the treadmill (i.e., training environment). We also quantified the steady-state before overground walking (**SteadyState**, average of last 40 strides) to get information about the final adapted state of each of the groups before testing participants overground. Then, we quantified early aftereffects overground (**EarlyPost**_**OG**_, averaged of first 5 strides) during the initial steps of the post-adaptation epoch to assess the generalization of movements from the treadmill (i.e., training environment) to overground walking (i.e., testing environment). We quantified the late aftereffects overground (**LatePost**_**OG**_, average of last 40 strides). This was done to verify that all participants returned to their baseline values overground before returning to the treadmill. Lastly, we looked at the magnitude of aftereffects on the treadmill (i.e., **EarlyPost**_**TM**_) to assess the remaining treadmill-specific motor patterns not washed out by walking overground. The data of one participant in the Young_Abrupt_ group during the treadmill post-adaptation epoch was not recorded due to technical difficulties. Therefore, this participant was excluded from the analysis of EarlyPost_TM_ only. We subtracted participant-specific biases on the treadmill or overground before aggregating the data of all individuals for group analyses. This was done by subtracting the baseline biases on the treadmill or overground that matched the specific walking condition. For example, we subtracted the Baseline bias of each participant measured on the treadmill from outcome measures recorded on the treadmill during adaptation and post-adaptation epochs.

Lastly, we quantified %Generalization, which is the magnitude of aftereffects overground for step length asymmetry SL_asym_ expressed as a percentage of treadmill aftereffects as shown below:

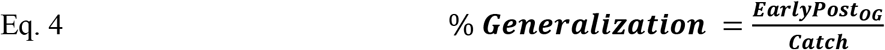

In this equation, EarlyPost_OG_ refers to the unbiased SL_asym_ values during early post-adaptation overground, and Catch refers to the unbiased SL_asym_ values during the catch period. A metric of %Generalization was not computed for other gait parameters because this was not numerically stable, resulting in unrealistic values of %Generalization for the Lead_asym_ and Trail_asym_ parameters. As an alternative, we performed a regression analysis to determine the relationship between the leading and trailing positions for each leg (*Lead*_*fast*_, *Lead*_*slow*_, *Trail*_*fast*_ and *Trail*_*slow*_) during post-adaptation on the treadmill (i.e., Catch) and overground (i.e., EarlyPost_OG_). This procedure is described in the statistical analysis section.

### 2.5 Statistical Analysis

#### 2.5.1 Planned Analysis

##### Power Analysis

The number of participants per group was determined using the overground aftereffects of step length asymmetry from old and young participants experiencing the same abrupt speed differences as in this study (Sombric et al., 2017). We specifically assumed the estimated error variance of 0.00097 and the estimated difference of 0.032 between the older and young groups (Sombric et al., 2017). We also anticipated a difference of 0.05 in the generalization between the gradual and abrupt groups based on a previous gradual vs. abrupt adaptation study (Torres-Oviedo and Bastian, 2012). This led to the effect size of 0.513 and 0.8 for older vs. young and gradual vs. abrupt comparisons, respectively. Our power analysis indicated that n = 8 participants per group would allow us to detect the anticipated difference between the older and young groups with at least 80% power and a significance level of 0.05. This sample size would also enable us to detect the expected difference between gradual and abrupt groups with 99% power. Therefore, we adopted a target sample size of 8 participants per group, which is comparable to the number of participants in other studies assessing the generalization of locomotor adaptation in older adults (Sombric et al., 2017; Sombric and Torres-Oviedo, 2021).

##### Group Analysis

We performed one-sample Kolmogorov–Smirnov tests to determine if each parameter (i.e., SL_asym_, Lead_asym_, and Trail_asym_) was normally distributed in every epoch of interest (i.e. Steady-StateBC, Catch, Steady-State, EarlyPost_OG_, LatePost_OG_, and EarlyPost_TM_) in all 4 groups. We found that all parameters were normally distributed, thus we ran separate two-way ANOVAs to test the effects of age (i.e., older vs. young) and perturbation schedule (i.e., gradual vs. abrupt) on each of our gait parameters. These two-way ANOVAs were performed on unbiased data (i.e., the condition-specific baseline was removed) to focus on changes that occurred beyond those due to distinct group biases. In case of a significant interaction effect, we performed post-hoc comparisons with Tukey corrections to identify differences between groups. A significance level of α = 0.05 was used for the two-way ANOVAs tests. Also, we wanted to determine if aftereffects were significant and participants go back to baseline behavior at the end of post-adaptation in each group. Therefore, we performed a one-sided one-sample t-test to determine whether Catch, EarlyPost_OG_, LatePost_OG_, and EarlyPost_TM_ values were different from zero. We corrected for multiple comparisons using a Benjamini–Hochberg procedure (Benjamini and Hochberg, 1995), as we have done before (Aucie et al., 2020), in which we corrected the significance threshold for each epoch by setting a false discovery rate of 5% (FDR correction). Consequently, a p-value < 0.044 was significant considering the FDR correction. Stata (StataCorp., Collage Station, TX, United States) was used to perform the ANOVAs and one-sample t-tests, whereas MATLAB (TheMathWorks, Inc., Natick, MA, United States) was used for all other analyses. P-values, F-values, and t-values are reported for all group analyses, whereas effect sizes (η^2^ for two-way ANOVAs, and Cohen’s d for unpaired t-test) were only reported when a significant effect size was found.

##### Individual Analysis

Previous studies have shown that speed-specific baseline values are predictive of steady-state behavior in the leading and trailing leg positions both in healthy (Sombric et al., 2019) and post-stroke survivors (Sombric and Torres-Oviedo, 2020). Therefore, we wanted to verify whether the same relationship holds in our data. Thus, we performed linear regression analysis for each group separately to quantify the similarity between lead and trail leg positions across speed-specific baseline and late adaptation epochs. To confirm the previous finding, we tested the model y = m*x, where y is the predicted leg position (e.g., Lead_fast_) during late adaptation and x is the leg position during baseline (Sombric et al., 2019; Sombric and Torres-Oviedo, 2020). These regressions were performed in data pooled by age or perturbation size if the group analysis revealed that either of these factors had a significant effect on the dependent variable (i.e., steady-state). For example, if we observed a significant age effect on the steady-states, we performed a regression per age group (e.g., Young_Abrupt_ & Young_Gradual_ pooled together and Old_Abrupt_ and Old_Gradual_ pooled together).

#### 2.5.2 Post-hoc Analysis

We unexpectedly observed that the maximum error size was not significantly different between the gradual and abrupt groups. In particular, gradual groups experienced errors, as large as those in the abrupt groups, during the initial steps of the re-adaptation condition following the catch condition. Therefore, we eliminated the catch condition in two additional groups (n=8 each) of older adults (10 males 6 females, mean age 76±5 years) adapted gradually (Old_gradual_NC_) or abruptly (Old_abrupt_NC_) without a catch. These participants simply experienced a resting break, rather than a catch condition. We compared the generalization between these two groups with significantly different error sizes upon gradual vs. abrupt adaptation (see Figure 2). More specifically, we used unpaired t-tests to compare the aftereffects between the two groups when either walking overground (EarlyPost_OG_, LatePost_OG_) or on the treadmill (EarlyPost_TM_). Of note, one participant in the Old_gradual_NC_ was excluded from the analysis because the ankle marker was not collected throughout the post-adaptation epochs due to technical difficulties. Also, some participants (i.e., n=3 in Old_abrupt_NC_ and n=5 in the Old_gradual_NC_) were not naïve to split-belt walking, but they had more than 6 weeks between experimental sessions, reducing the potential effect of split-belt exposure on overground aftereffects (Sombric et al., 2017).

**Figure 2.**
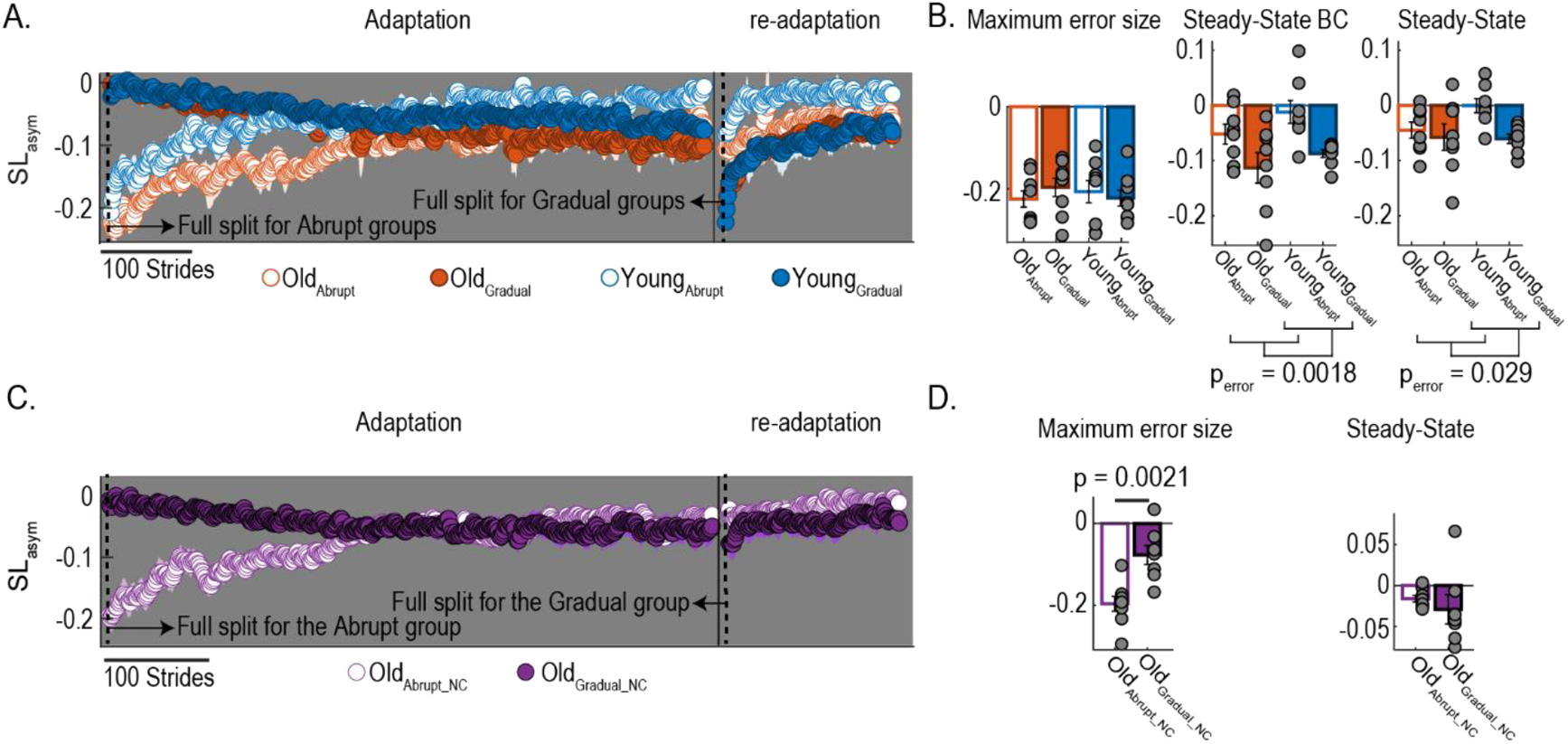
Time courses of step length asymmetry (i.e., performance error) during the adaptation and re-adaptation periods. **(A, C)** Time courses of step length asymmetry for all groups. Colored dots represent the group average of five consecutive strides and colored shaded regions indicate the standard error for each group. The full split time point used to compute the maximum error size, for each group is indicated by the black dashed line. (**B, D**) Bar plots indicate the mean and standard errors for each group for the error size and steady-states. Gray dots represent values for individual participants. Note that the reported values are unbiased (i.e., bias in each participant during baseline on the treadmill was subtracted before calculating outcome measures).

## 3 Results

### Gradual adaptation led to small errors only in the absence of a catch period

We observed that participants in the gradual and abrupt groups had the same error size (maximum errors) throughout the split-belt condition. We compared the error size when the belts had reached the same speeds (i.e. 0.5 m/s for the slow belt and 1 m/s for the fast belt) in both adaptation conditions. We found that the error size was the same in all groups. This was indicated by no significant effects of age (F_age(1,28)_ = 0.03, p_age_ = 0.86), perturbation schedule (F_error(1,28)_ = 0.07, p_error_ = 0.79), or interaction (F_interaction(1,28)_ = 0.85, p_interaction_ = 0.36) on step length asymmetry SL_asym_ (Figure 2B). Only participants in the older gradual group without the catch (Old_gradual_NC_; Figure 2C; filled purple dots) experienced a smaller error size during adaptation compared to the older abrupt groups without the catch (Old_abrupt_NC_; Figure 2C; empty purple dots). This was indicated by the significant maximum error size difference between these groups (t = −3.83, p = 0.0021, d = −1.98) (Figure 2D). Thus, the gradual groups experienced smaller errors compared to the abrupt groups only in the absence of the catch period.

### Large errors led to a higher steady-state during adaptation in all age groups

In general, we observed a significant effect of adaptation condition on the steady-state that people reached during split-belt walking. In other words, older and young participants corrected their step length asymmetry more when adapted abruptly compared to when adapted gradually. Figure 2A indicates the time course for SL_asym_ during adaptation and re-adaptation. Note that older and young participants adapted abruptly (i.e., empty dots) reached a steady-state closer to zero compared to those adapted gradually (i.e., filled dots) at the end of the adaptation (SteadyStateBC) and re-adaptation periods (SteadyState). Accordingly, there was a significant effect of adaptation condition on steady-state before the catch (F_error(1,28)_ = 11.92, p_error_ = 0.0018, η^2^ = 0.29) and at the end of re-adaptation period (F_error(1,28)_ = 5.34, p_error_ = 0.0285, η^2^ = 0.16) (Figure 2B). Steady-states were not different between the groups in the absence of a catch period (t = 0.76; p = 0.46), which was driven by an outlier data point (Figure 2D). Specifically, we found a significantly lower steady-state for the gradual group without the catch (filled purple dots) compared to the abrupt without the catch (empty purple dots) (t = 3.02; p = 0.011; d = 1.07) when this outlier individual is excluded from the analysis. Lastly, we did not find any age or interaction effect for SL_asym_ at steady-state before the catch (F_age(1,28)_ = 2.79, p_age_ = 0.11; F_interaction(1,28)_ = 0.13, p_interaction_ = 0.72) or at the end of re-adaptation period (F_age(1,28)_ = 1.74, p_age_ = 0.19; F_interaction(1,28)_ = 2.17, p_interaction_ = 0.15). In summary, the steady-state values of SL_asym_ were not affected by the participant’s age but depended on the perturbation schedule during split-belt walking in the presence of a catch condition.

Furthermore, we observed that the steady-state differences in SL_asym_ across groups were driven by the asymmetry in the Leading legs (i.e., Lead_asym_), but not by the asymmetry of trailing legs (i.e., Trail_asym_). Figure 3A shows that Lead_asym_ for older and young participants adapted gradually reached a smaller adapted state (Figure 3A; filled dots) than those adapted abruptly (Figure 3A; empty dots) at the end of both adaptation and re-adaptation periods (before and after the catch). On the other hand, Trail_asym_ reached a similar adapted state across groups. Accordingly the Lead_asym_ at steady-state before the catch (F_error(1,28)_ = 14.03, p_error_ = 0.0008, η^2^ = 0.33) and at the end of adaptation (F_error(1,28)_ = 5.81, p_error_ = 0.023, η^2^ = 0.17) was larger for older and young groups adapted gradually than those adapted abruptly (Figure 3B). For the Trail_asym_ we observed an effect of adaptation type and age on the steady-state before catch (F_age(1,28)_ = 4.22, p_age_ = 0.049, η^2^ = 0.13; F_error(1,28)_ = 5.32, p_error_ = 0.028, η^2^ = 0.16), but these effects go away by the end of the re-adaptation period (F_age(1,28)_ = 1.74, p_age_ = 0.19; F_error(1,28)_ = 1.56, p_error_ = 0.22) (Figure 3D). Similar results were observed in the steady states of Lead_asym_ and Trail_asym_ when gradual and abrupt groups were adapted without a catch (Lead_asym_: t = 1.17; p = 0.26; Trail_asym_: t = −0.14; p = 0.89, data not shown). Therefore, all participants reached a similar trailing asymmetry at the end of the adaptation period, whereas the leading asymmetry was smaller at steady-state in the abrupt than gradual groups.

**Figure 3.**
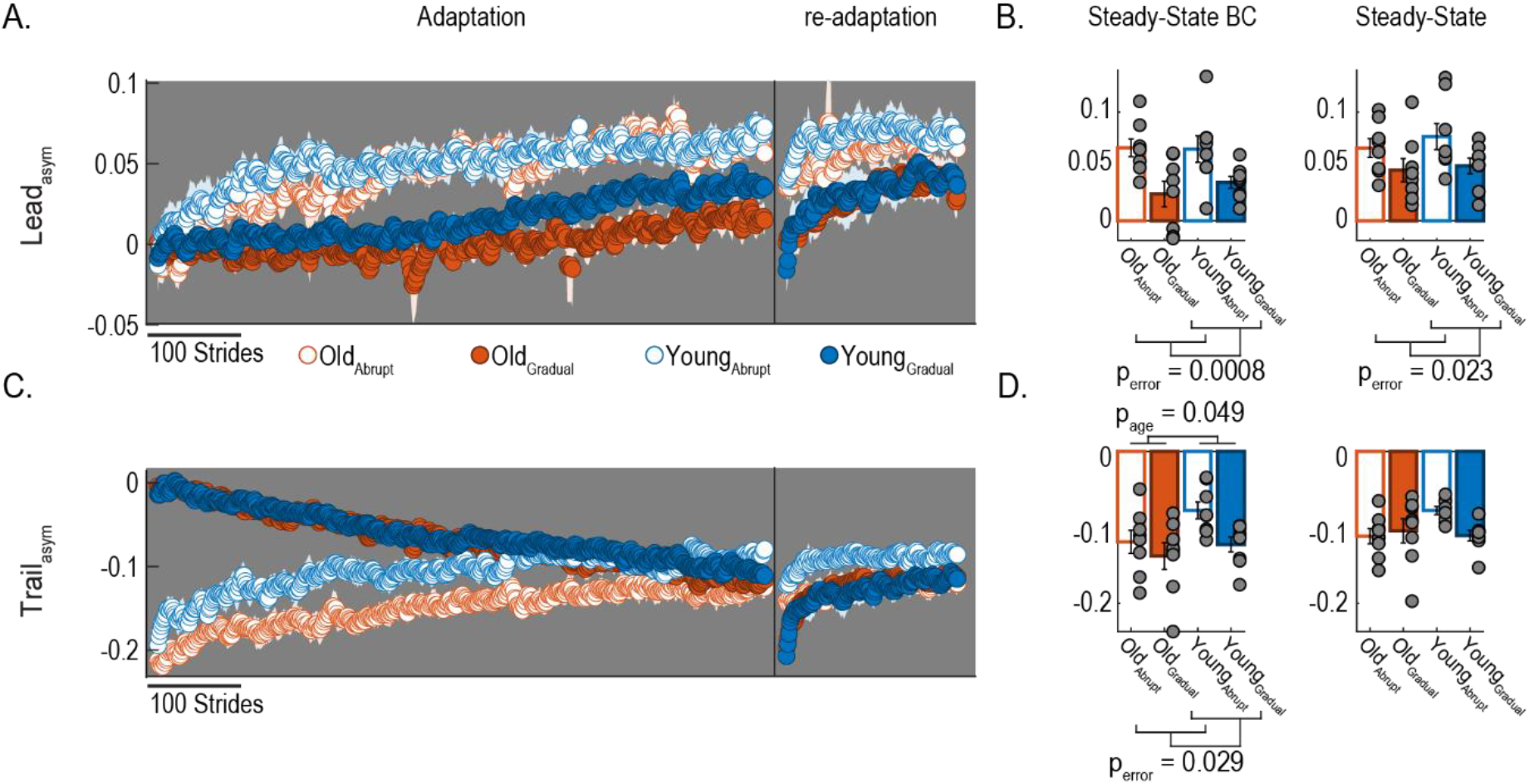
Adaptation of lead and trail asymmetries. **(A, C)** Time courses for lead and trail asymmetry during adaptation. Colored dots represent the group average of five consecutive strides and colored shaded regions indicate the standard error for each group. (**B, D**) Bar plots indicate the mean and standard errors for each group for steady-state before (i.e., Steady-State BC) and after (i.e., Steady-State) the catch. Gray dots represent individual participants. Note that the reported values are unbiased. This was done by subtracting the bias in each participant during baseline walking on the treadmill at medium speed (i.e., 1 m/s).

### Baseline leg’s positions predict the steady states before and after the catch in all age groups when adapted gradually or abruptly

It has been shown recently that participants recover the baseline speed-specific, leading and trailing, leg’s position during steady-state split-belt walking (Sombric et al. 2019; Sombric and Torres-Oviedo 2020). We found that this relation between baseline and steady-state split-belt walking was not altered by age or perturbation size. This was indicated by the significant relationship between the speed-specific baseline and steady-states found in each of the four groups (Old_Abrupt_, Old_Gradual,_ Young_Abrupt_, and Young_Gradual_) before (R^2^> 0.63; p < 0.001) and after the catch (R^2^> 0.59; p < 0.001). Also, our group analysis indicated that the perturbation scheduled affected the steady-state values of SL_asym_. Thus, we grouped the participants by how they were adapted (Abruptly vs. Gradually). We found that the speed-specific baseline values were predictor of steady-state behavior both before (Abrupt: R^2^ = 0.86; p < 0.0001 SS = 1.01*speed-specific baseline; Gradual: R^2^ = 0.82; p < 0.0001 SS = 0.98*speed-specific baseline) and after the catch (Abrupt: R^2^ = 0.87; p < 0.0001 SS = 1.01*speed-specific baseline; Gradual: R^2^ = 0.86; p < 0.0001 SS = 0.98*speed-specific baseline) as shown in Figure 4. In sum, the steady-state movements in participants of all ages are strongly correlated to their baseline behavior, regardless of how the split-belt perturbation is introduced during the adaptation period.

**Figure 4.**
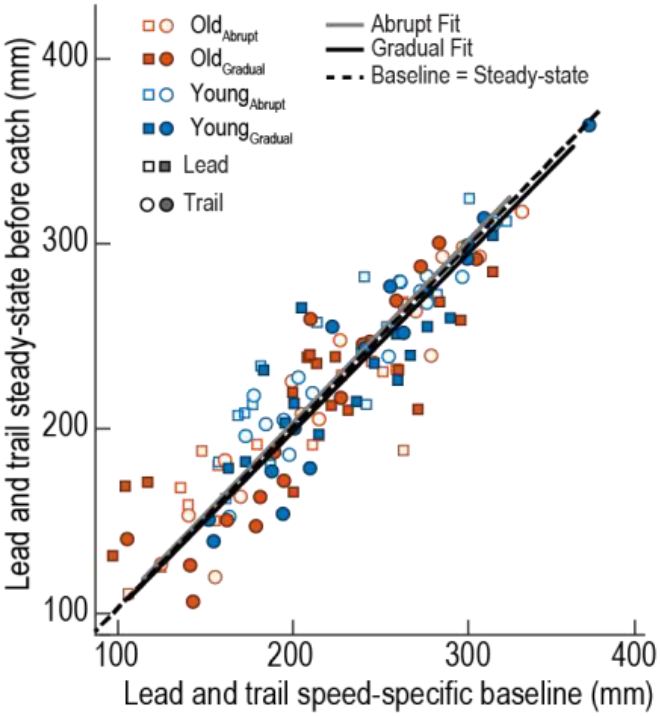
The similarity between lead and trailing leg’s positions across speed-specific baseline and steady-state before catch epochs is illustrated by the significant regression of abrupt (y = m*x, 95% Confidence interval for m = [0.98, 1.04]) and gradual fits (y = m*x, 95% Confidence interval for m = [0.95, 1.01]). Note that the regression lines closely overlap with the idealized situation (dashed line) in which baseline and steady-state values are identical (dashed line; slope of one, i.e., y = x).

### Abrupt perturbations result in greater aftereffects on the treadmill in young and older age groups

Participants adapted gradually had smaller sensorimotor recalibration compared to those adapted abruptly regardless of age. This is indicated by the significantly smaller aftereffects during the catch period in the gradual (Fig. 5A, filled circles) than abrupt groups (Fig. 5A, empty circles). Figure 5A shows the averaged aftereffects during the catch condition on the treadmill for older and young participants adapted gradually or abruptly. Error size had a significant effect on the aftereffects during the catch for SL_asym_ (F_error(1,28)_ = 20.31, p_error_ = 0.0001, η^2^ = 0.42), Lead_asym_ (F_error(1,28)_ = 14.27, p=0.0008, η^2^ = 0.34), and Trail_asym_ (F_error(1,28)_ = 15.45, p=0.0005, η^2^ = 0.36). On the other hand, we did not have an age or interaction effect in any of the parameters during the catch (SL_asym_: F_age(1,28)_ = 3.72, p_age_ = 0.064; F_interaction(1,28)_ = 0.38, p_interaction_ = 0.54; Lead_asym_: F_age(1,28)_ = 2.45, p_age_ = 0.13; F_interaction(1,28)_ = 0.64, p_interaction_ = 0.43; Trail_asym_: F_age(1,28)_ = 2.99, p_age_ = 0.095; F_interaction(1,28)_ = 0.07, p_interaction_ = 0.79). Moreover, we observed a significant treadmill aftereffect for SL_asym_ and Trail_asym_ in all the groups (See Table 1), but not in Lead_asym_ for the Old_gradual_ group, which is marginally significant (p = 0.057, t = 2.28) (Figure 5B). Overall, we observed that the perturbation schedule during adaptation, but not the participants’ age, altered the aftereffects on the treadmill.

**Table 1.** Aftereffects during catch on the treadmill (Catch)

| Table 1. Aftereffects during catch on the treadmill (Catch) |  |  |  |
| --- | --- | --- | --- |
| Outcome measure for each Group | p-value | t-value | Effect size (Cohen's d) |
| SL<sub>asym</sub> |  |  |  |
| Old<sub>Abrupt</sub> | <0.0001 | 10.9 | 3.89 |
| Old<sub>Gradual</sub> | 0.0003 | 6.76 | 2.39 |
| Young<sub>Abrupt</sub> | <0.0001 | 11.7 | 4.14 |
| Young<sub>Gradual</sub> | 0.0005 | 6.18 | 2.18 |
| Lead<sub>asym</sub> |  |  |  |
| Old<sub>Abrupt</sub> | 0.0035 | 4.31 | 1.52 |
| Old<sub>Gradual</sub> | 0.057 (ns) | 2.28 | N/A |
| Young<sub>Abrupt</sub> | 0.0001 | 7.72 | 2.73 |
| Young<sub>Gradual</sub> | 0.034 | 2.64 | 0.93 |
| Trail<sub>asym</sub> |  |  |  |
| Old<sub>Abrupt</sub> | <0.0001 | 19.3 | 6.82 |
| Old<sub>Gradual</sub> | <0.0001 | 11.85 | 4.19 |
| Young<sub>Abrupt</sub> | <0.0001 | 12.33 | 4.36 |
| Young<sub>Gradual</sub> | 0.0002 | 6.97 | 2.47 |

**Figure 5.**
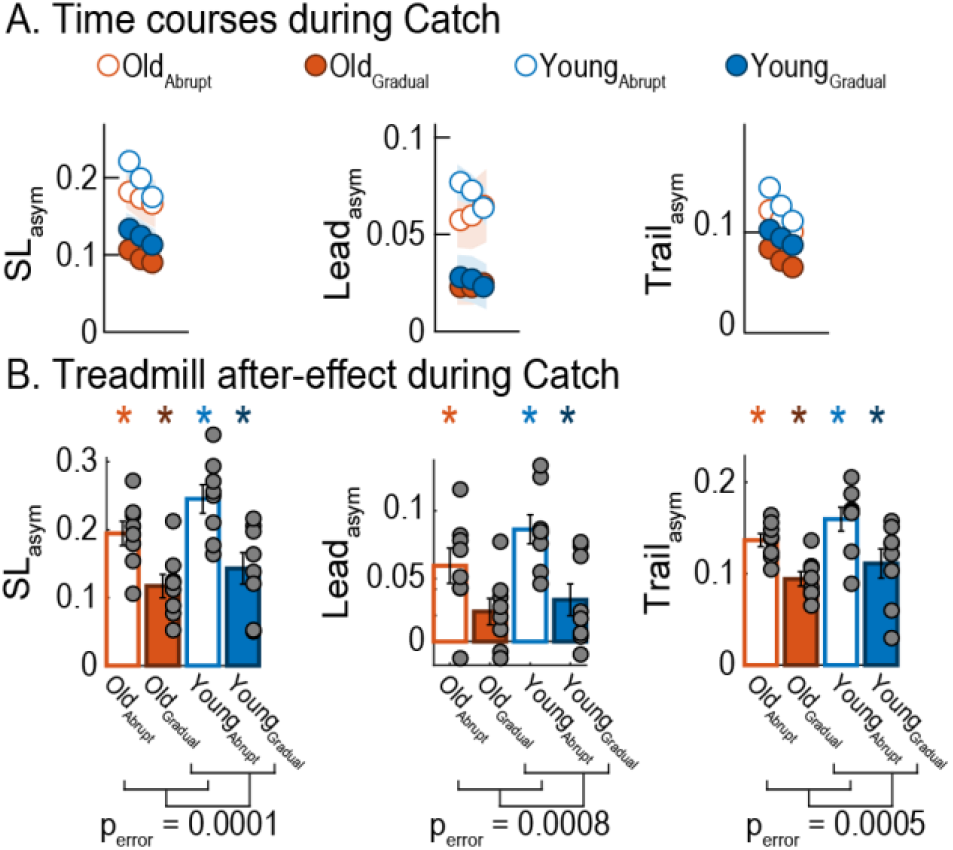
Treadmill aftereffect (i.e., Catch) for asymmetry parameters. **(A)** Time courses for step length asymmetry, lead asymmetry, and trail asymmetry during the catch condition. Colored dots represent the group average of five consecutive strides and colored shaded regions indicate the standard error for each group. (**B**) Bar plots indicate the mean and standard errors for each group for the catch (i.e., Catch) condition. Gray dots represent individual participants. Note that the reported values are unbiased. This was done by subtracting the bias in each participant during baseline walking on the treadmill at medium speed (i.e., 1 m/s).

### Older participants generalized more than young when adapted gradually or abruptly

We found that age had a significant effect on the generalization of locomotor adaptation. Figure 6 shows timecourses of SL_asym_ during overground post-adaptation in groups experiencing a catch (Figure 6A) and those who did not (Figure 6B). While all groups exhibited initial aftereffects (i.e., EarlyPost_OG_) that were significantly different from zero (See EarlyPost_OG_ results in Table 2), older adults (i.e., orange and purple dots) have larger aftereffects when walking overground than young groups (i.e., blue dots). Accordingly, we found a significant effect of age on the initial overground aftereffects (i.e., EarlyPost_OG_) for SL_asym_ in the groups experiencing a catch (F_age(1,28)_ = 10.68, p_age_ = 0.0029, η^2^ = 0.28) (Figure 6C; Old_Abrupt_, Old_Gradual,_ Young_Abrupt_, and Young_Gradual_). This age effect was also observed in %Generalization of SL_asym_ (F_age(1,28)_ = 7.6, p_age_ = 0.01, η^2^ = 0.21), which quantifies the initial aftereffects overground (testing context) as a percentage of the initial aftereffects on the treadmill (training context).

**Table 2.** Initial aftereffects during overground post-adaptation (EarlyPostOG)

|  |  |  |  |
| --- | --- | --- | --- |
| **Outcome measure for each Group** | **p-value** | **t-value** | **Effect size (Cohen's d)** |
| **SL<sub>asym</sub>** |  |  |  |
| Old<sub>Abrupt</sub> | 0.0039 | 4.24 | 1.49 |
| Old<sub>Gradual</sub> | <0.0001 | 10.2 | 3.62 |
| Young<sub>Abrupt</sub> | 0.0001 | 7.43 | 2.63 |
| Young<sub>Gradual</sub> | 0.013 | 3.33 | 1.18 |
| Old<sub>Abrupt_NC</sub> | 0.0039 | 4.23 | 1.49 |
| Old<sub>Gradual_NC</sub> | 0.0009 | 6.02 | 2.28 |
| **Lead<sub>asym</sub>** |  |  |  |
| Old<sub>Abrupt</sub> | 0.066 (ns) | 2.18 | N/A |
| Old<sub>Gradual</sub> | 0.088 (ns) | 1.98 | N/A |
| Young<sub>Abrupt</sub> | 0.53 (ns) | 0.66 | N/A |
| Young<sub>Gradual</sub> | 0.60 (ns) | -0.55 | N/A |
| Old<sub>Abrupt_NC</sub> | 0.14 (ns) | 1.65 | N/A |
| Old<sub>Gradual_NC</sub> | 0.035 | 2.72 | 1.03 |
| **Trail<sub>asym</sub>** |  |  |  |
| Old<sub>Abrupt</sub> | 0.0008 | 5.65 | 1.99 |
| Old<sub>Gradual</sub> | <0.0001 | 11.15 | 3.94 |
| Young<sub>Abrupt</sub> | 0.0001 | 7.41 | 2.62 |
| Young<sub>Gradual</sub> | 0.0017 | 4.94 | 1.75 |
| Old<sub>Abrupt_NC</sub> | 0.0003 | 6.54 | 2.31 |
| Old<sub>Gradual_NC</sub> | 0.0015 | 5.52 | 2.09 |
| **Final aftereffects during overground post-adaptation (LatePost<sub>OG</sub>)** |  |  |  |
| **Outcome measure for each Group** | **p-value** | **t-value** | **Effect size (Cohen's d)** |
| **SL<sub>asym</sub>** |  |  |  |
| Old<sub>Abrupt</sub> | 0.0006 | 5.83 | 2.06 |
| Old<sub>Gradual</sub> | 0.0022 | 4.72 | 1.67 |
| Young<sub>Abrupt</sub> | 0.0027 | 4.53 | 1.6 |
| Young<sub>Gradual</sub> | 0.024 | 2.87 | 1.01 |
| Old<sub>Abrupt_NC</sub> | 0.022 | 2.94 | 1.04 |
| Old<sub>Gradual_NC</sub> | 0.014 | 3.42 | 1.29 |
| **Lead<sub>asym</sub>** |  |  |  |
| Old<sub>Abrupt</sub> | 0.0013 | 5.2 | 1.84 |
| Old<sub>Gradual</sub> | 0.013 | 3.32 | 1.17 |
| Young<sub>Abrupt</sub> | 0.004 | 4.21 | 1.49 |
| Young<sub>Gradual</sub> | 0.044 | 2.46 | 0.87 |
| Old<sub>Abrupt_NC</sub> | 0.0821 (ns) | 2.03 | N/A |
| Old<sub>Gradual_NC</sub> | 0.0061 | 4.14 | 1.56 |
| **Trail<sub>asym</sub>** |  |  |  |
| OldAbrupt | 0.0014 | 5.09 | 1.79 |
| OldGradual | 0.0046 | 4.09 | 1.45 |
| YoungAbrupt | 0.018 | 3.07 | 1.09 |
| YoungGradual | 0.015 | 3.20 | 1.13 |
| OldAbrupt_NC | 0.012 | 3.35 | 1.19 |
| OldGradual_NC | 0.032 | 2.78 | 1.05 |

**Figure 6.**
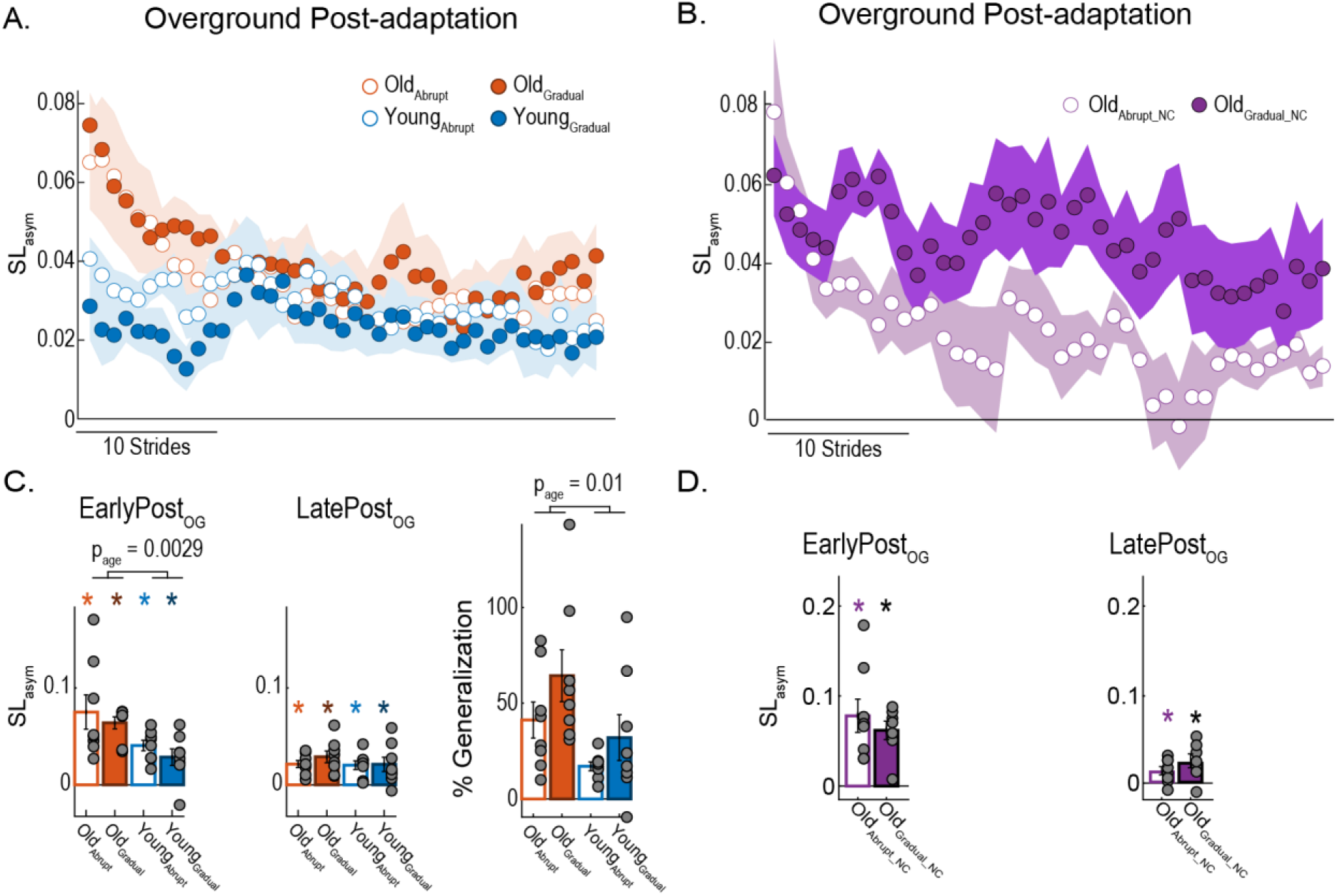
Generalization of step length asymmetry. (A-B) Time courses for step length asymmetry during overground walking post-adaptation for groups experiencing a catch condition (panel A) and groups without a catch (panel B). Colored dots represent the group average of five consecutive strides and colored shaded regions indicate the standard error for each group. (C-D) Bar plots indicate the mean and standard errors for each group’s initial (EarlyPost_OG_) and final (LatePost_OG_) aftereffects during overground post-adaptation. Group aftereffects are displayed for the groups experiencing a catch condition (panel C) and groups without a catch (panel D). Asterisks denote group averages significantly different from zero. (E) %Generalization for each group experiencing a catch condition. This measure indicates the size of initial aftereffects expressed as a percentage of the aftereffects on the treadmill during the catch. In all, gray dots represent individual participants. Note that the reported values are unbiased. This was done by subtracting the bias in each participant during baseline walking overground.

Conversely, perturbation schedule did not impact the generalization of SL_asym_ quantified as raw values (F_error(1,28)_ = 1.17, p_error_ = 0.29; F_interaction(1,28)_ < 0.001, p_interaction_ = 0.97) or as a %Generalization (F_error(1,28)_ = 3.46, p_error_ = 0.73; F_interaction(1,28)_ = 0.16, p_interaction_ = 0.69). This finding was unexpected given previous reports indicating that gradual adaptation results in more overground aftereffects compared to abrupt adaptation (Torres-Oviedo and Bastian, 2012; Alcantara et al.2018). We considered that the equally large errors between our gradual and abrupt groups experiencing a catch could explain the similar generalization across these perturbation schedules. Thus, we tested two additional groups of older adults who adapted gradually or abruptly without a catch (Old_Abrupt_NC_ and Old_Gradual_NC_). The generalization patterns of these additional groups confirmed our finding that error size did not have an impact on the generalization of older adults. While these groups had significantly different error sizes during adaptation (Figure 2), they exhibit similar initial overground aftereffects (t = 0.73; p = 0.48) (Figure 6D). In sum, the participants’ age, but not the perturbation schedule during adaptation, regulated the generalization of movements from the treadmill to overground.

The age-mediated differences in overground aftereffects vanished by the end of the post-adaptation period overground (i.e., LatePost_OG_). Namely, we found that old and young groups maintained aftereffects that were significantly different from zero by the end of the post-adaptation period overground (See LatePost_OG_ results in Table 2). However, neither age, perturbation schedule, nor an interaction between these two factors had a significant effect on any of the asymmetry parameters at the end of the post-adaptation period (SL_asym_: F_age(1,28)_ = 0.62, p_age_ = 0.44; F_error(1,28)_ = 0.54, p_error_ = 0.47; F_interaction(1,28)_ = 0.34, p_interaction_ = 0.56; Lead_asym_: F_age(1,28)_ = 0.33, p_age_ = 0.57; F_error(1,28)_ < 0.001, p_error_ = 0.95; F_interaction(1,28)_ = 0.02, p_interaction_ = 0.9; Trail_asym_: F_age(1,28)_ = 0.59, p_age_ = 0.45; F_error(1,28)_ = 1.15, p_error_ = 0.29; F_interaction(1,28)_ = 0.64, p_interaction_ = 0.43) (Figure 6C). Similarly, SL_asym_ values at the end of overground post-adaptation were similar between the older adults adapted abruptly or gradually without a catch (Old_Abrupt_NC_ vs. Old_Gradual_NC_; t = −1.44; p = 0.17) (Figure 6D). Therefore, while participants did not fully return to their baseline asymmetry values by the end of the post-adaptation period overground, neither age or perturbation schedule had an impact on the remaining aftereffects.

Similar results were observed in the other asymmetry measures: age had a significant effect on generalization of adapted movements, but perturbation type did not. All groups exhibited initial overground aftereffects that were significantly different from zero in Trail_asym_ but not in Lead_asym_ (See EarlyPost_OG_ results in Table 2). Only the older participants adapted gradually without a catch (Old_Gradual_NC_ group) had significant overground aftereffects in all asymmetry measures including Lead_asym_ (p = 0.035, t = 2.72, d = 1.03) (data not shown). Time courses are shown in figure 7A and 7B. We observed that older adults (i.e., orange dots) have larger aftereffects when walking overground than young groups (i.e., blue dots) in Trail_asym_ but this effect was much smaller in Lead_asym_. Accordingly, we found a significant effect of age on the initial overground aftereffects (i.e., EarlyPost_OG_) of Trail_asym_ (F_age(1,28)_ = 9.53, p_age_ = 0.0045, η^2^ = 0.25) and Lead_asym_, but the effect size was smaller in the latter one (F_age(1,28)_ = 5.63, p_age_ = 0.025, η^2^ = 0.17). Interestingly, perturbation schedule did not impact the generalization of aftereffects in any parameter (Lead_asym_: F_error(1,28)_ = 1.38, p_error_ = 0.25; F_interaction(1,28)_ = 0.07, p_interaction_ = 0.79; Trail_asym_: F_error(1,28)_ = 0.48, p_error_ = 0.49; F_interaction(1,28)_ = 0.09, p_interaction_ = 0.77) contrasting our anticipated results. The age-mediated differences in overground aftereffects was not observed by the end of the post-adaptation period overground (i.e., LatePost_OG_). Namely, all groups exhibited significant aftereffects at the end of the post-adaptation period overground (See LatePost_OG_ results in Table 2), except for Old_Abrupt_NC_ group (p = 0.0821, t = 2.03) in Lead_asym_ (data not shown). However, neither age, perturbation schedule, nor an interaction between these two factors had a significant effect on the final aftereffects (LatePost_OG_) for both asymmetry measures (Lead_asym_: F_age(1,28)_ = 0.33, p_age_ = 0.57; F_error(1,28)_ < 0.001, p_error_ = 0.95; F_interaction(1,28)_ = 0.02, p_interaction_ = 0.9; Trail_asym_: F_age(1,28)_ = 0.59, p_age_ = 0.45; F_error(1,28)_ = 1.15, p_error_ = 0.29; F_interaction(1,28)_ = 0.64, p_interaction_ = 0.43). In conclusion, older adults adapted gradually or abruptly generalized more than young the asymmetric pattern from the treadmill to overground walking.

**Figure 7.**
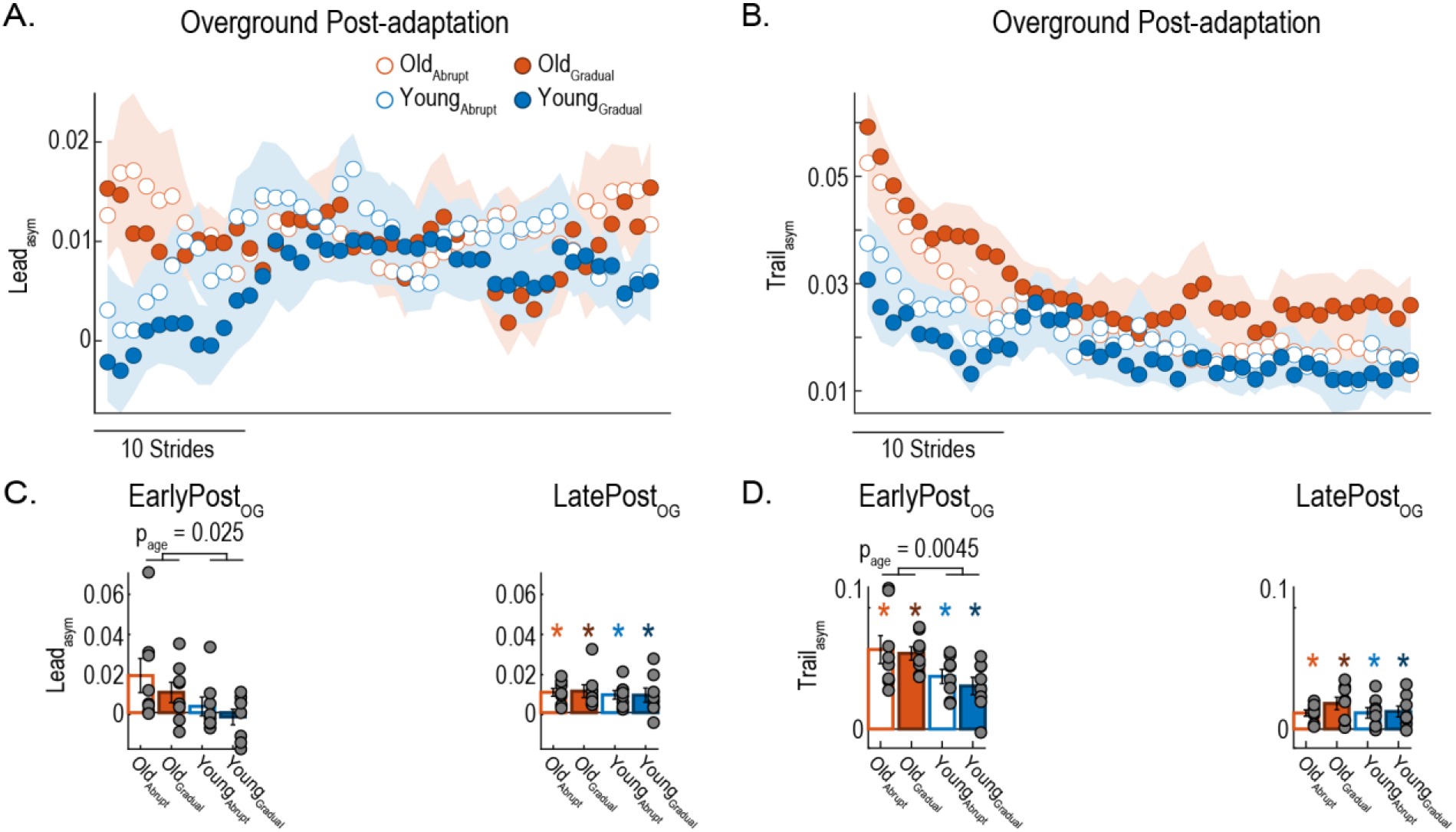
Overground aftereffects of lead asymmetry and trail asymmetry. (A-B) Time courses of the aftereffects in lead asymmetry and trail asymmetry during overground post-adaptation. Colored dots represent the group average of five consecutive strides and colored shaded regions indicate the standard error for each group. (**C-D**) Bar plots indicate the mean and standard errors for each group for the initial aftereffects (i.e., EarlyPost_OG_) and final aftereffects (i.e., LatePost_OG_) during the overground post-adaptation. Gray dots represent individual participants. Asterisks denote group averages significantly different from zero. Note that the reported values are unbiased. This was done by subtracting the bias in each participant during baseline walking overground.

### Overground walking washes out more the treadmill aftereffects in the gradual than in the abrupt groups

We observed that the perturbation scheduled affected the remaining aftereffects when participants returned to the treadmill after walking overground. Figure 8A, B, C, and D show the time courses of the remaining aftereffects during post-adaptation on the treadmill for all of the groups. We observed that all groups had aftereffects that were significantly different from zero in all asymmetry parameters (see Table 3). This was expected since participants did not return to their baseline gait before they walked again on the treadmill. However, young and older adults who adapted gradually (filled circles) had smaller remaining aftereffects than those who adapted abruptly (empty circles). This is clearly observed in the older adult groups adapted without a catch (purple dots). Consistently, the initial aftereffects during treadmill post-adaptation is significantly smaller in the Old_gradual_NC_ than in the Old_abrupt_NC_ groups (t = 4.02; p = 0.0015, d=2.08). The groups adapted with a catch show a similar tendency. However, the effect of perturbation condition was only significant in the Lead_asym_ (F_error(1,27)_ = 4.68, p_error_ = 0.039, η^2^ = 0.15) and not in SL_asym_ (F_error(1,27)_ = 2.29, p_error_ = 0.14) or Trail_asym_ (F_error(1,27)_ = 0.57, p_error_ = 0.46). We also found that neither age (SL_asym_: F_age(1,27)_ = 0.39, p_age_ = 0.54; Lead_asym_: F_age(1,27)_ = 0.03, p_age_ = 0.87; Trail_asym_: F_age(1,27)_ = 0.88, p_age_ = 0.36) nor an interaction between age and perturbation schedule significantly modified the washout of treadmill aftereffects by overground walking (SL_asym_: F_interaction(1,27)_ = 0.04, p_interaction_ = 0.84; Lead_asym_: F_interaction(1,27)_ = 0.01, p_interaction_ = 0.93; Trail_asym_: F_interaction(1,27)_ = 0.08, p_interaction_ = 0.78). In sum, overground walking washed out more the treadmill aftereffects when participants were adapted gradually than when they were adapted abruptly.

**Table 3.** Initial aftereffects during treadmill post-adaptation (EarlyPostTM)

|  |  |  |  |
| --- | --- | --- | --- |
| **Outcome measure for each Group** | **p-value** | **t-value** | **Effect size (Cohen's d)** |
| **SL<sub>asym</sub>** |  |  |  |
| Old<sub>Abrupt</sub> | 0.0006 | 5.86 | 2.07 |
| Old<sub>Gradual</sub> | 0.0001 | 7.97 | 2.82 |
| Young<sub>Abrupt</sub> | 0.0003 | 7.49 | 2.83 |
| Young<sub>Gradual</sub> | 0.0002 | 7.23 | 2.56 |
| Old<sub>Abrupt_NC</sub> | <0.0001 | 12.9 | 4.56 |
| Old<sub>Gradual_NC</sub> | <0.0001 | 12.1 | 4.56 |
| **Lead<sub>asym</sub>** |  |  |  |
| Old<sub>Abrupt</sub> | 0.0059 | 3.90 | 1.38 |
| Old<sub>Gradual</sub> | 0.0034 | 4.34 | 1.53 |
| Young<sub>Abrupt</sub> | 0.0005 | 6.89 | 2.61 |
| Young<sub>Gradual</sub> | 0.0083 | 3.64 | 1.29 |
| Old<sub>Abrupt_NC</sub> | 0.0006 | 5.92 | 2.09 |
| Old<sub>Gradual_NC</sub> | 0.0332 | 2.75 | 1.04 |
| **Trail<sub>asym</sub>** |  |  |  |
| Old<sub>Abrupt</sub> | 0.0002 | 7.38 | 2.61 |
| Old<sub>Gradual</sub> | <0.0001 | 11.01 | 3.89 |
| Young<sub>Abrupt</sub> | 0.0003 | 7.55 | 2.85 |
| Young<sub>Gradual</sub> | 0.0001 | 8.27 | 2.92 |
| Old<sub>Abrupt_NC</sub> | <0.0001 | 20.4 | 7.21 |
| Old<sub>Gradual_NC</sub> | <0.0001 | 11.68 | 4.41 |

**Figure 8.**
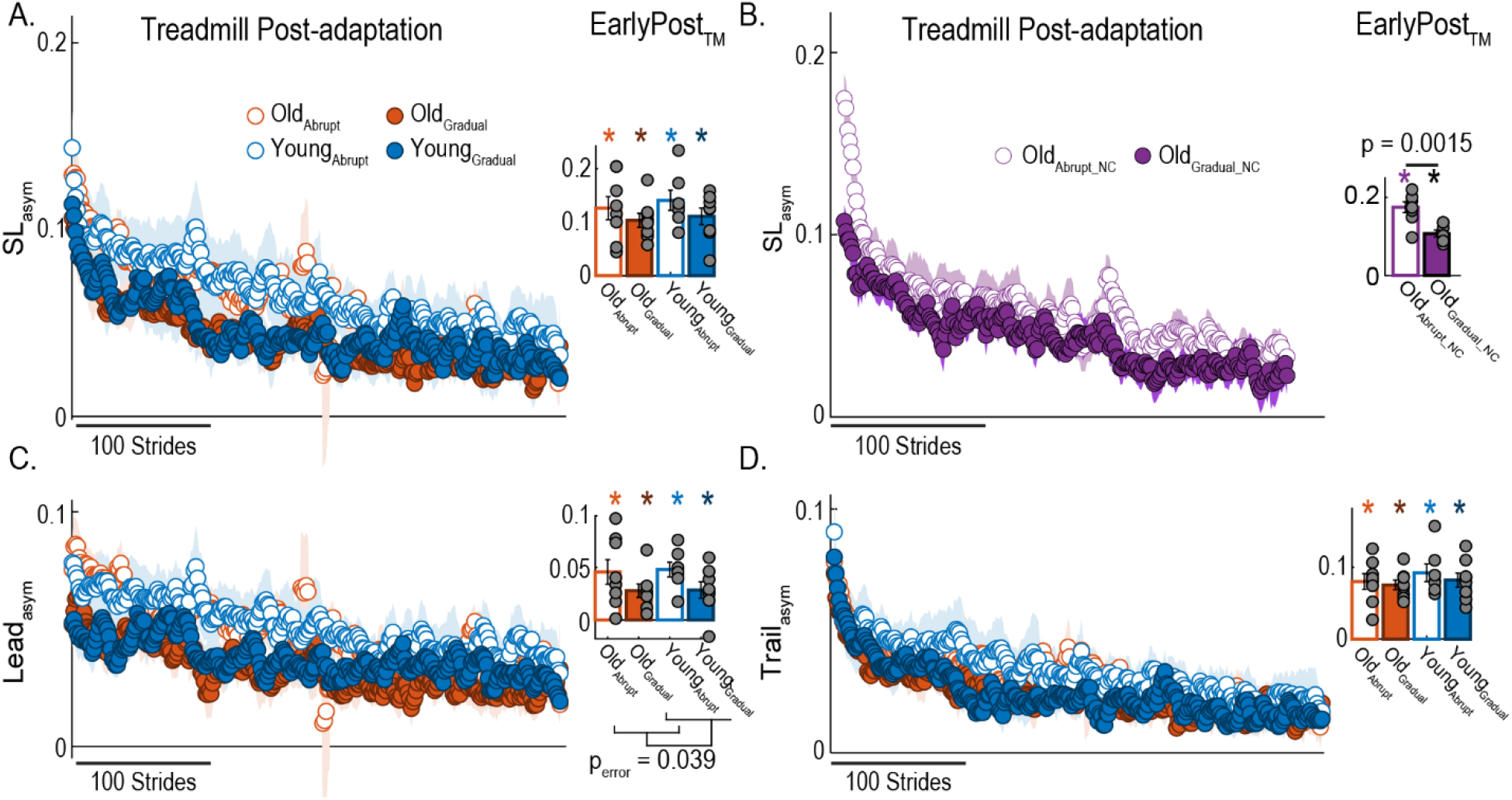
Aftereffects on the treadmill of all asymmetry parameters. (A-D - left side) Time courses for step length asymmetry when walking on the treadmill during post-adaptation for groups experiencing a catch condition (panel A) and groups without a catch (panel B). Time courses for lead asymmetry (panel C) and trail asymmetry (panel D) for the groups experiencing a catch are also shown. Colored dots represent the group average of five consecutive strides and colored shaded regions indicate the standard error for each group. (A-D - right side) Bar plots indicate the mean and standard errors for each group’s initial aftereffects on the treadmill following overground walking. Gray dots represent individual participants. Asterisks denote group averages significantly different from zero. Note that the reported values are unbiased. This was done by subtracting the bias in each participant during baseline walking on the treadmill at medium speed (i.e., 1 m/s).

## 4 Discussion

We investigated the interaction between age and split-perturbation size, which are two factors known to influence the generalization of locomotor adaptation. We found that experiencing abrupt adaptation (large split-perturbation) led to greater steady states and aftereffects on the treadmill compared to gradual adaptation (small split-perturbation) in both age groups. On the other hand, we observed that older adults generalized their movements to overground walking more than young regardless of the perturbation size during adaptation. Our regression analysis revealed that the movements during baseline at different speeds predict the behavior of steady states for both age groups. Lastly, we found that the overground walking following adaptation washes out the effect of gradual adaptation more than abrupt. Taken together our results suggest that older adults generalize more than young regardless of the error size experienced during adaptation.

### Large errors led to higher steady-states and aftereffects on the treadmill in both age groups

Participants in the abrupt groups reached a higher steady-state compared to the gradual groups during the adaptation at the presence of catch in both young and older age groups. This means that large errors led to more adaptation of movements compared to small errors when the catch was present. On the other hand, in our post-hoc analysis (i.e., no catch trial), we found that both abrupt and gradual groups had the same steady-states. This finding is aligned with the fact that gradual adaptation leads to fragile motor memories (Roemmich and Bastian, 2015) and it is more susceptible to washout by the catch trial as observed in our groups with the catch. In other words, experiencing catch is similar to experiencing the perturbation in the opposite direction that could washout the fragile motor memories in the gradual groups. Another explanation is that the participants in the gradual groups with the catch experienced large errors during the re-adaptation phase as if they were adapted for the first time (Roemmich and Bastian, 2015); however, they adapted for a shorter period compared to the abrupt adaptation (300 strides vs. 900 strides). We know from reaching studies that a short exposure of the abrupt perturbation may not be enough to fully adapt the movement (Joiner et al., 2013). Therefore, it is possible that the re-adaptation period of 300 strides was not long enough to fully recover the adapted states in gradual groups with the catch, while in the case of no catch, participants got exposed to the same perturbation without any disruption, which helped them store the recalibrated movements easier.

We also observed that the participants in the abrupt groups have higher aftereffects on the treadmill (i.e., more sensorimotor recalibration) compared to the participants in the gradual groups in both young and older age groups. In other words, large errors led to more sensorimotor recalibration of movements within the same environment compared to small errors, which is consistent with previous findings (Torres-Oviedo and Bastian, 2012). Other locomotor adaptation studies have shown that an abrupt but not gradual change in the walking environment engages neural processes to acquire and store explicit knowledge about the new environment (Roemmich and Bastian, 2015). This fact can explain why the abrupt adaptation facilitated the storage of the recalibrated movements compared to the gradual adaptation (i.e., small errors). Moreover, the storage of the recalibrated movements is slower with the gradual than the abrupt perturbation (Taylor et al., 2014); therefore, our gradual groups may not be fully adapted by the end of the adaptation phase. Another explanation for the fact that the gradual and the abrupt perturbation led to different aftereffects, can be that they result in different internal representations of the learning environment (Roemmich and Bastian, 2015). For instance, in reaching literature, it has been shown that the adaptation to an abrupt perturbation is driven by feedforward motor planning, whereas adaptation to a gradual perturbation is dependent on feedback control (Saijo and Gomi, 2010).

While the perturbation schedule during adaptation altered the steady states and aftereffects on the treadmill, we did not find any effect of age in these epochs. This observation is consistent with previous studies showing that sensorimotor recalibration upon external perturbations is not impaired in older adults for walking (Bruijn et al., 2012; Malone and Bastian, 2016; Sombric et al., 2017) and reaching behaviors (Buch et al., 2003; Bock, 2005; Bock and Girgenrath, 2006) for the kinematic metrics. In contrast, our finding is not consistent with the previous literature on muscle activity. Iturralde and Torres-Oviedo’s model showed that as age increases the adaptability of movements decreases (Iturralde and Torres-Oviedo, 2019) in the muscle domain. This suggests that the control of limb movements is affected by age at the muscle domain; however, those changes are not translated to the kinematic level of our movements as shown by our data. Taken together, the error size experienced during the adaptation plays a significant role in sensorimotor adaptation and recalibration while the age of participants is only affecting the muscle and not the kinematic domains.

### Older participants generalized more than young in gradual and abrupt perturbations

We found that age had a significant effect on the generalization of locomotor adaptation, regardless of the error size experienced during adaptation. Previous studies have shown that older adults have a larger transfer of adapted movements across conditions in reaching (Fernández-Ruiz et al., 2000; Bock and Girgenrath, 2006; Heuer and Hegele, 2008) and walking (Sombric et al., 2017), which is consistent with our findings. In other words, older adults exhibit greater motor perseveration compared to younger adults when they switch across different environments. There are three potential explanations for these age-related differences. First, more motor perseveration in older adults can be explained by the degeneration of basal ganglia as we age. We know that the basal ganglia are responsible for motor switching (Brown and Almeida, 2011; Leunissen et al., 2013; Balser et al., 2014). This means that the age-related structural (Wolpe et al., 2020) and functional (Bäckman et al., 2006; Ota et al., 2006; Walhovd et al., 2011) changes in the basal ganglia can lead to poor motor switching and greater motor perseveration. Second, older adults might rely more on the movements that they have just learned during adaptation. Because healthy aging results in higher motor (Kallio et al., 2012; Vanden Noven et al., 2014) and sensory noise (Zhang et al., 2008; Goble et al., 2009; Maheu et al., 2015); therefore, they have less sensitivity to errors. This means that older adults try to generate the same pattern that they were doing at the end of adaptation instead of updating their movement (i.e., deadapting the learned movements) as they transition to the overground walking condition. Lastly, older adults are naturally more variable in their movements compared to young adults (Osoba et al., 2019). This means that older adults tend to attribute the sensed errors during adaptation to their own faulty movements (Berniker and Kording, 2008; Kelly and Sober, 2014); therefore, having difficulty switching their motor patterns across different conditions (Sombric et al., 2017; Sombric and Torres-Oviedo, 2021). Taken together, our results showed that older adults have a harder time switching and disengaging their motor patterns when facing a new environment compared to young adults.

On the other hand, we found that error size does not affect the generalization of movements from the treadmill (i.e., training environment) to the overground (i.e., testing environment). This is at odds with previous literature showing that gradual adaptation leads to more generalization than abrupt adaptation in both young adults (Torres-Oviedo and Bastian, 2012) and post-stroke patients (Alcântara et al., 2018). This discrepancy might be because our gradual groups did not adapt as much as our abrupt groups, which had not been reported before (Torres-Oviedo and Bastian, 2012; Alcântara et al., 2018). The reduced adaptation of our gradual groups compared to the abrupt is indicated by the lower steady-states, which is known to be positively correlated to less sensorimotor recalibration (i.e., less aftereffects on the treadmill) (Sombric et al., 2019; Aucie et al., 2020). This deficit in the adaptation of gradual groups compared to abrupt groups might have not been observed before because we used a smaller perturbation size (speed difference) than the one used before (Torres-Oviedo and Bastian, 2012; Alcântara et al., 2018). Smaller perturbation sizes are known to induce less adaptation and smaller aftereffects (Finley et al., 2015; Morehead et al., 2015; Marinovic et al., 2017; Yokoyama et al., 2018). Our interpretation is consistent with previous work showing that gradual adaptation results into more fragile motor memories than abrupt adaptation in the absence of an extended period of adaptation at the fully perturbed state (Roemmich and Bastian, 2015). Lastly, smaller aftereffects indicate less sensorimotor recalibration, but also make it more challenging to detect between group differences in generalization, which is a metric with large inter-subject variability. Taken together, our results suggest that the generalization of aftereffects beyond the training environment might be limited by the extent to which people adapt their movements in said training environment.

Another factor reducing the generalization of the gradual groups compared to the abrupt groups in those with the catch trial might have been the large asymmetries induce by the catch trial itself. Notably, the gradual groups experiencing the catch trial exhibited the same maximum error size as the abrupt groups before overground post-adaptation. Thus, participants in the gradual groups with the catch trial did not truly experience smaller errors than those in the abrupt groups. These large errors in the gradual groups could have been perceived as out-of-the ordinary, limiting the generalization of movements (Torres-Oviedo and Bastian, 2012). The catch trial could have also limited the generalization of movements because removing the perturbation during the catch is a perturbation in-and-of itself (Herzfeld et al., 2014; Iturralde and Torres-Oviedo, 2019), which is repeated when people walk overground. Thus, the repeated exposure to removing the split-perturbation might facilitate switching between distinct walking patterns within the training environment or across distinct walking environments. We are currently testing this hypothesis in another study explicitly assessing the impact of catch trials on the generalization of movements.

### Speed specific baseline positions are a predictor of steady-state before and after the catch regardless of the error size

We found that the speed-specific baseline values are a predictor of steady-state behavior both before and after the catch for both gradual and abrupt groups despite the differences during the steady states. This finding is consistent with previous literature showing that all participants recover their baseline leg orientation in both young healthy adults (Sombric et al., 2019) and post-stroke survivors (Sombric and Torres-Oviedo, 2020). We know that leg orientation is closely regulated to walk at distinct speeds (Orendurff et al., 2008). Therefore, participants need to take speed-specific step lengths to adapt to the split-belt environment. In other words, participants would match the speed-specific step length (i.e., biomechanically driven) by late adaptation regardless of their age or the error size that they are experiencing during the adaptation. This implies that neither participants’ age nor the error size during adaptation can impact the biomechanical constrain set by the speed of the belts when walking on the treadmill.

While all groups recovered their baseline leg orientation at the steady-state at the individual level, we found that the error size impacted the steady states that participants reach in leading, but not trailing asymmetry measures regardless of age groups. We know from previous studies that the lead and trail asymmetry are correlated to spatial and temporal features of the gait, respectively (Mariscal et al., 2020; Sombric and Torres-Oviedo, 2020). Therefore, differences in the adaptation of lead and trail asymmetry can be explained by the fact that the spatial and temporal features of the gait adaptation are controlled by different neural substrates (Malone and Bastian, 2014) in post-stroke patients. This has also been reported in other split-belt paradigms (Bruijn et al., 2012; Vervoort et al., 2019) in older adults. In summary, our findings confirmed that the spatial and temporal gait features adapt differently, but are constrained to the speed at which the participant is walking at.

### Overground walking washes out more the treadmill aftereffects in the gradual than in the abrupt groups

We found that overground walking washed out more the treadmill aftereffects when participants were adapted gradually than when they were adapted abruptly. Specifically, we observed this in Lead_asym_ for the groups that experienced the catch trial. This is in odds with previous literature reporting no differences between the gradual and abrupt adaptation in the spatial domain (Lead_asym_ is correlated to the spatial domain) when the participants return to the training environment (Torres-Oviedo and Bastian, 2012). One interpretation is that overground walking only washed out the aftereffect in SL_asym_ and Trail_asym_ because these were the only adapted movement that carried over to overground walking and consequently, that was washout by overground walking. We did not observe any difference in the SL_asym_ and Trail_asym_. This observation was consistent with the previous literature showing no differences between the gradual and abrupt adaptation in SL_asym_ and temporal (Trial_asym_ is correlated to the temporal domain) measures(Torres-Oviedo and Bastian, 2012; Alcântara et al., 2018). On the other hand, we found that in the groups without the catch there is a difference between the gradual and the abrupt group even in SL_asym_. This means that gradual adaptation leads to fragile motor memories (Roemmich and Bastian, 2015), that is, motor memories that are susceptible to washout by walking in other contexts, which is particularly clear in the people walking without a catch. Another interpretation is that participants adapted more abruptly than gradually (Torres-Oviedo and Bastian, 2012; Roemmich and Bastian, 2015), thus, the remaining aftereffects have to be greater for the abrupt than the gradual condition for the same extent of washout by the overground experience. To sum up, gradual adaptation leads to motor memories that are more susceptible to washout by overground walking, and this susceptibility increases by introducing disruption during adaptation.

### Clinical Implications

Our results confirmed that older adults have difficulties switching motor patterns (Fernández-Ruiz et al., 2000; Bock and Girgenrath, 2006; Heuer and Hegele, 2008; Sombric et al., 2017; Sombric and Torres-Oviedo, 2021), which leads to a higher risk of falls (Lockhart et al., 2002). Thus, the risk of falling in the older population could decrease by practicing switching between various walking terrains (Wagner et al., 1994; Tinetti et al., 1996). We also showed that healthy aging leads to larger generalization regardless of the error size experienced during adaptation. This is particularly important because our findings suggest that large errors might be better than small when training older populations. Specifically, we demonstrated that older adults learn more from large than small errors, and large errors do not limit the generalization of this learning.

Therefore, the older clinical population will carry over the enhanced learning from large errors to real-life situations. Finally, the age-related differences in generalization also suggest that older patients have a higher chance of motor improvements beyond the clinical setting compared to younger ones. In conclusion, our work highlights the importance of age-related changes in the generalization of locomotor adaptation, which could be used to improve rehabilitation techniques beyond clinical settings.

## 5 Author’s contribution statement

YA contributions include analysis, and interpretation of the data, drafting the work and agreement to be accountable for all aspects of the work. HMH and CJS contributions include acquisition of data and processing. GT contributions include conception and design of the work, revising the work and agreement to be accountable for all aspects of the work. All authors contributed to revising the manuscript and providing a final approval of the version to be published.

## 7 Acknowledgments

We thank Dr. Patrick Sparto for his assistance in recruitment of participants in this study.

